# Haploidy-linked disruption of tissue organization in zebrafish larval eyes

**DOI:** 10.64898/2026.11.10.765796

**Authors:** Daiki Saito, Ayano Suzuki, Kan Yaguchi, Riho Sakai, Yuki Yokoi, Kiminori Nakamura, Shigenobu Yonemura, Ryota Uehara

## Abstract

In vertebrate species, somatic haploidy generally causes embryonic or larval lethality with drastic morphological defects. However, the detrimental effects of haploidy at the cellular or tissue level remain largely unknown. In this study, we described cellular and tissue defects associated with haploidy in zebrafish larval eyes as a tissue model system. The haploid larval retina showed overall disorganized tissue layer structures. Photoreceptor cells, which usually align in the outermost tissue layer, were reduced in number and severely displaced, with drastic retardation of their connecting cilia and outer segments specifically in haploids. Ganglion cells in the inner retinal layer were also reduced in abundance and abnormally organized in haploids. These multifaceted cellular and tissue defects indicate that haploid eyes suffer severe visual functional defects, in addition to well-recognized size reductions. Our findings provide insight into diverse cellular principles that may limit the developmental capability of vertebrates in the haploid state.

**Summary statement:** Cross-ploidy comparison of zebrafish larval eyes revealed haploidy-specific defects in lamination and retinal cellular organization.

## Introduction

Diplontic life cycles are common among a wide range of animal groups. In vertebrates, haploid embryos, arising through irregular activation of unfertilized oocytes, cannot achieve full development (Komen and Thorgaard, 2007; Ocalewicz, 2024; Wutz, 2014). Although haploidy intolerance is a stringent constraint on the evolution of animal life cycles, its underlying principles remain largely unknown. Recent years, cross-ploidy comparative studies in culture cells have revealed pleiotropic defects occurring in association with haploidy, such as chronic centrosome loss (Yaguchi et al., 2018; Yoshizawa et al., 2020), mitotic instability (Olbrich et al., 2017; Yaguchi et al., 2018), delay in DNA replication (Edwards et al., 2021), and organelle dysfunction (Di Minin et al., 2026; Ishida-Ishihara et al., 2024). In contrast, cross-ploidy comparisons at multicellular tissue levels remain limited, precluding the establishment of organismal principles of haploidy intolerance.

In mammals, haploidy-linked embryonic lethality arises at the pre-implantation stage, accompanied by drastic cell-division defects (Yaguchi et al., 2018), precluding investigation of haploidy-linked effects on organogenesis. In contrast, haploid embryos often survive through organogenesis in non-mammalian vertebrates, such as frogs and fish, making them ideal models for studying haploid intolerance in multicellular processes. For example, earlier studies in haploid amphibian or fish embryos have revealed morphologically detectable, gross organ malformations in haploids, which are collectively referred to as haploid syndromes. More recently, cross-ploidy comparisons using live imaging in zebrafish embryos have revealed different aspects of haploidy-linked problems in multicellular reorganization, such as organ growth retardation associated with mitotic abnormalities (Yaguchi et al., 2024) or impaired collective movement during gastrulation (Menon et al., 2020). Meanwhile, it remains largely unknown whether and how haploidy affects higher-order tissue organization, governed by diverse cell types, during organogenesis.

This study systematically investigated the influence of haploidy on multicellular organization in the zebrafish eye at ultrastructural, intracellular, and tissue levels. We chose the eye as a model because it is an excellent system for studying multi-tissue coordination and diverse cell function (Amini et al., 2022; Norden, 2023). The zebrafish eye is particularly useful for imaging analyses because of its transparency, making it a widely used organ research system (Fadool and Dowling, 2008; Richardson et al., 2017). Moreover, the retardation of eye growth has long been recognized as a hallmark of haploid syndromes in non-mammalian vertebrates (Luo and Li, 2003; Yaguchi et al., 2024). However, descriptions of eye defects in haploid embryos have been limited to general morphological observations (Luo and Li, 2003; Purdom, 1969), and it remained unclear which cell types or tissues are influenced by haploidy. Here, we found pleiotropic problems caused by haploidy in different biological layers, including cell differentiation, intracellular structure formation, extracellular matrix organization, and multi-tissue rearrangements. Our findings add new entries to the catalog of haploidy-affected bioprocesses, providing insights into the organismal principles underlying ploidy constraints in animals.

## Results

### Drastic tissue disorganization in haploid larval eyes

To understand the effects of haploidy on multicellular tissue organization, we first analyzed the overall structures of eyes isolated from 5-days post-fertilization (dpf) haploid or diploid zebrafish larvae, obtained through in vitro fertilization with UV-irradiated or non-irradiated sperm, respectively (Yaguchi et al., 2024), using electron microscopy (EM; Fig. 1A; see Materials and Methods). Diploid eyes showed highly ordered retinal tissue layers: The ganglion cell layer (GCL), the inner plexiform layer (IPL), the inner nuclear layer (INL), the outer plexiform layer (OPL), and the photoreceptor cell layer (PCL) formed in order from the innermost layer outward, arranged in a smooth semicircle surrounding the lens (Fig. 1A) (Amini et al., 2018; Schmitt and Dowling, 1999). In haploid eyes, the retinal layers frequently lost their smooth concentric arrangement around the lens, resulting in locally distorted tissue architecture accompanied by discontinuous IPL and OPL (Fig. 1A). The degree of disruption of the layered structure varied among haploid larvae. In severe cases, individual retinal layers became highly uneven in thickness. Moreover, the IPL and OPL were drastically retarded, and retinal layer boundaries were often difficult to trace continuously across the tissue (Fig. 1A). Consistent with the EM observation, fluorescence microscopy of nuclei and F-actin (stained by DAPI and fluorescent phalloidin) also showed drastic disorganization of retinal layer structures, with the F-actin-enriched IPL and OPL highly distorted and waving in haploid eyes (Fig. 1C). Therefore, haploid larval eyes were not only small in size but also exhibited profound abnormalities in their tissue organization.

**Figure 1:**
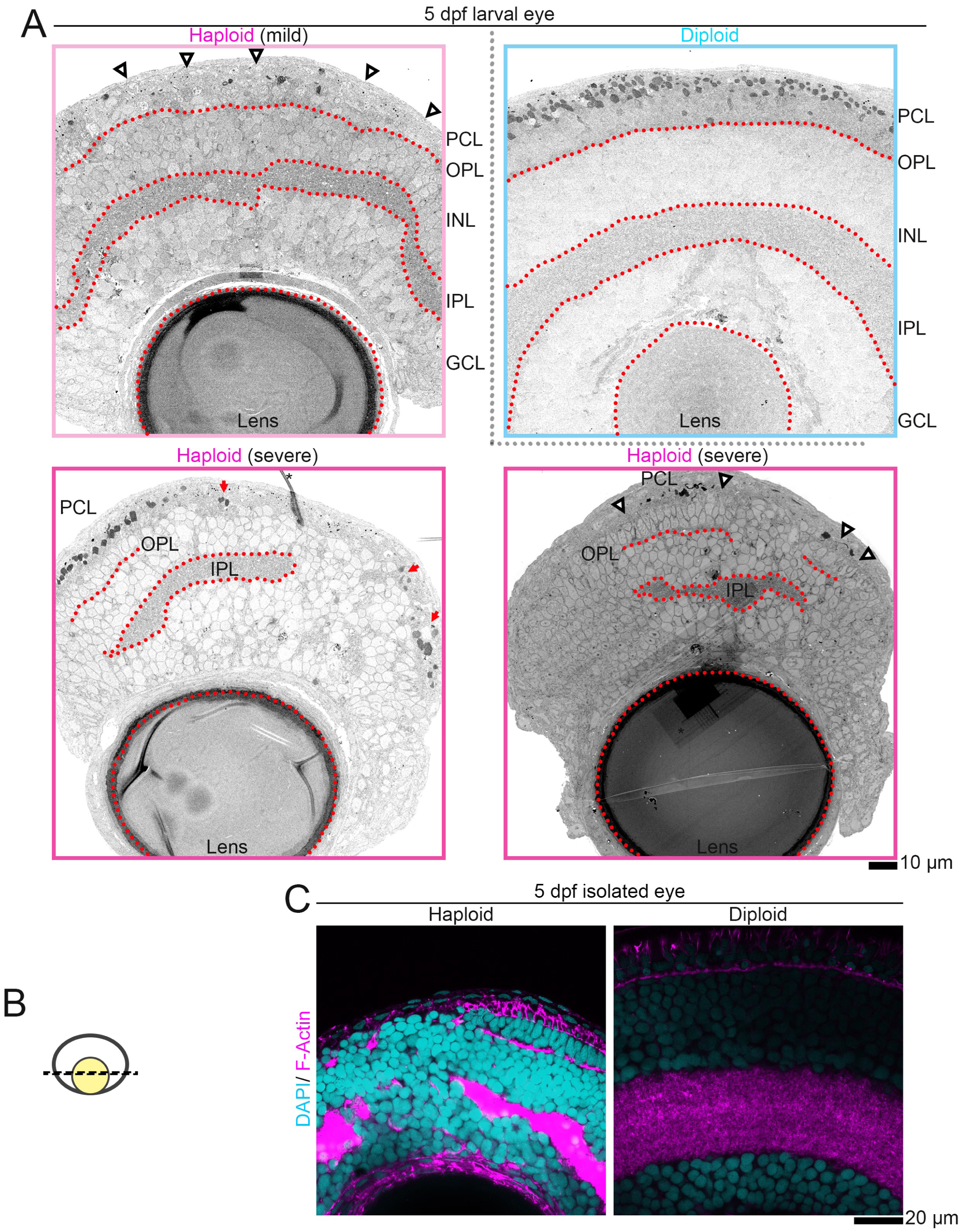
Haploidy-linked defects in retinal layer tissue structures. **(A)** EM images of retinal layer structures in eyes isolated from 5-dpf haploid or diploid larvae. The ganglion cell layer (GCL), the inner plexiform layer (IPL), the inner nuclear layer (INL), the outer plexiform layer (OPL), and the photoreceptor cell layer (PCL) are delineated by broken lines. Arrowheads indicate the PCL area devoid of outer segments of photoreceptor cells. Arrows indicate positions where the layer structure is disrupted, yet the outer segments of photoreceptor cells remain. The asterisk indicates artificial debris. Scale bar: 10 μm. **(B)** The scheme showing the location of the incision and the area observed within the eye. **(C)** Fluorescence microscopy of DNA and F-actin in isolated eyes from 5-dpf haploid or diploid larvae. Scale bar: 20 μm. Representative data from 5 eyes from 2 independent experiments.

### Defects in morphology and alignment of photoreceptor cells in haploids

In the EM images of the haploid retina above, photoreceptor cells, characterized by their electron-dense outer segments, were sparsely distributed and often separated by unusually large gaps along the outer retinal edge (Fig. 1A and S1). As a result, the PCL was only partially occupied by photoreceptor cells in many haploid eye regions (Fig. 1A and S1). This pattern sharply contrasted with the diploid retina, where photoreceptor cells were densely aligned throughout the PCL. Therefore, we further analyzed photoreceptor cell distributions by immunostaining for Arrestin3 (a photoreceptor cell marker) and acetylated α-tubulin (a connecting cilia marker that also marks neurons) in isolated haploid or diploid eyes at 5 dpf. In diploid retina, Arrestin3-marked photoreceptor cells were regularly aligned throughout the PCL with their connecting cilia extended toward the retinal periphery (Fig. 2A). Consistent with EM observation, photoreceptor cells were distributed in an intermittent manner in the PCL in haploids, with their substantial proportion mislocated as isolated cells deep within the neural retina (Fig. 2A). Proportion of Arrestin3-positive photoreceptor cells found in the outer edge of the PCL (per total photoreceptor cells in a confocal slice) was drastically reduced in haploids (29.9%) than in diploids (99.9%), verifying the above trend (Fig. 2B). Total number of Arrestin3-positive cells per single confocal eye section also significantly reduced in haploids than in diploids (Fig. 2C). Consistently, proportion of cells in the outer edge of neural retina that possessed connecting cilia (per total outer edge cells) was also significantly reduced in haploids (29.8%) than in diploids (97.7%; Fig. 2D). These data demonstrate that haploids suffer severe defects in both photoreceptor cell formation and PCL organization.

**Figure 2:**
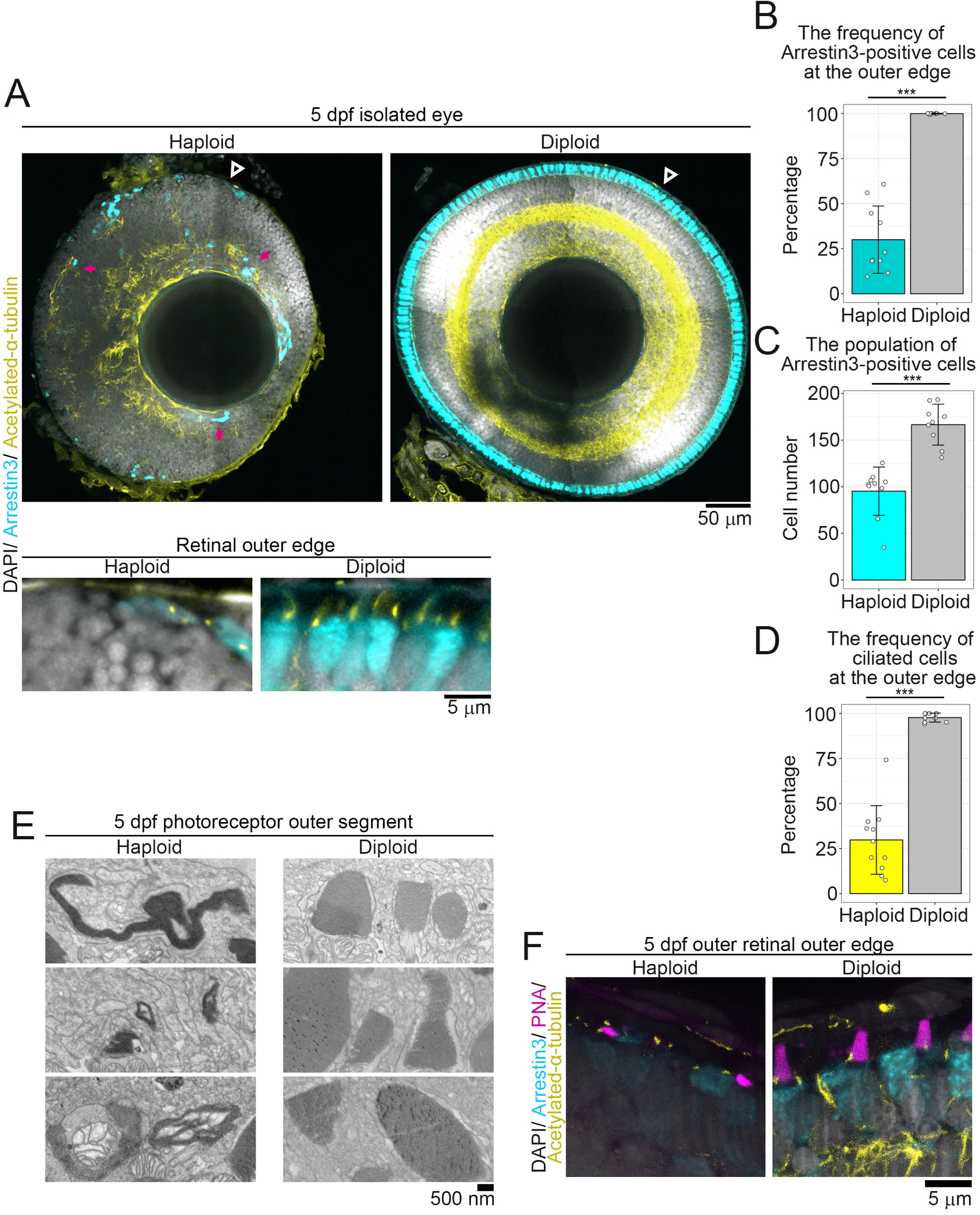
Haploidy-linked malformation of photoreceptor cells. **(A)** Immunostaining of Arrestin3 and acetylated-α-tubulin in eyes isolated from 5-dpf haploid or diploid larvae. DNA was stained with DAPI. Magenta arrowheads indicate Arrestin3-positive photoreceptor cells ectopically located in inner retinal tissue layers. Top: whole-eye view. Scale bar: 50 μm. Bottom: enlarged view of the retinal outer edge indicated by black arrowheads in the top images. Scale bar: 5 μm. **(B)** Frequency of Arrestin3-positive cells at the retinal outer edge, among all Arrestin3-positive cells, within single confocal eye sections. **(C)** Number of Arrestin3-positive cells within single confocal eye sections. **(D)** Frequency of ciliated cells, among all cells at the retinal outer edge, within single confocal eye sections. Mean ± standard deviation (s.d.) of at least 9 eyes from at least 5 larvae in two independent experiments, both for haploids and diploids. At least 35, 35, or 29 cells per eye were analyzed in B, C, or D, respectively, for each condition. ***p<0.001, two-tailed t-test). **(E)** EM images of photoreceptor outer segments in 5-dpf haploid or diploid larval retina. Scale bar: 500 nm. Representative data from 3 eyes from 2 independent experiments. **(G)** Fluorescence microscopy of PNA-labelled photoreceptor outer segments in 5-dpf haploid or diploid larval retina. Arrestin3 and acetylated-α-tubulin were co-immunostained, recapitulating the haploidy-specific severe retardation of outer segments in E. Scale bar: 5 μm.Representative data from 4 eyes from 2 independent experiments.

To further investigate potential defects in photoreceptor cell formation, we examined outer-segment morphology in high-magnification EM images. Compared to the well-organized membrane layers within the smoothly elongated outer segments in diploid photoreceptor cells, outer segments in haploids were severely deformed, showing bent, sinuous, invaginated, or inflected morphologies (Fig. 2E). In addition to membrane deformation, haploid outer segments frequently lost their uniform longitudinal orientation (Fig. 2E). The malformed outer segments were also observed with fluorescence-labeled peanut agglutinin (PNA), which specifically stained cone outer segment-associated matrix (Lessieur et al., 2017; Shi et al., 2017) (Fig. 2F). Three-dimensional reconstituted images of the PCL revealed that, not only outer segments, but also entire photoreceptor cells were often mis-oriented, consistent with the irregularity of the PCL organization in haploids (Supplemental Video S1, S2).

We next examined the morphology of connecting cilia in photoreceptor cells, which are important for outer-segment organization and photoreceptor function (Beryozkin et al., 2021; Hanke-Gogokhia et al., 2018; Krock et al., 2009; Ojeda Naharros et al., 2018). We observed connecting cilia with their basal bodies (marked by acetylated α-tubulin and γ-tubulin immunostaining, respectively) in the subretinal space of retinas using super-resolution imaging (Fig. 3A). The average length of connecting cilia significantly decreased in haploids (1.7 ± 0.8 μm), compared to diploids (2.4 ± 0.7 μm; Fig. 3B), with a less uniform size distribution. These observations collectively demonstrate that haploidy leads to abnormalities in photoreceptor cell architecture, at least with respect to connecting cilia length control and outer segment organization.

**Figure 3:**
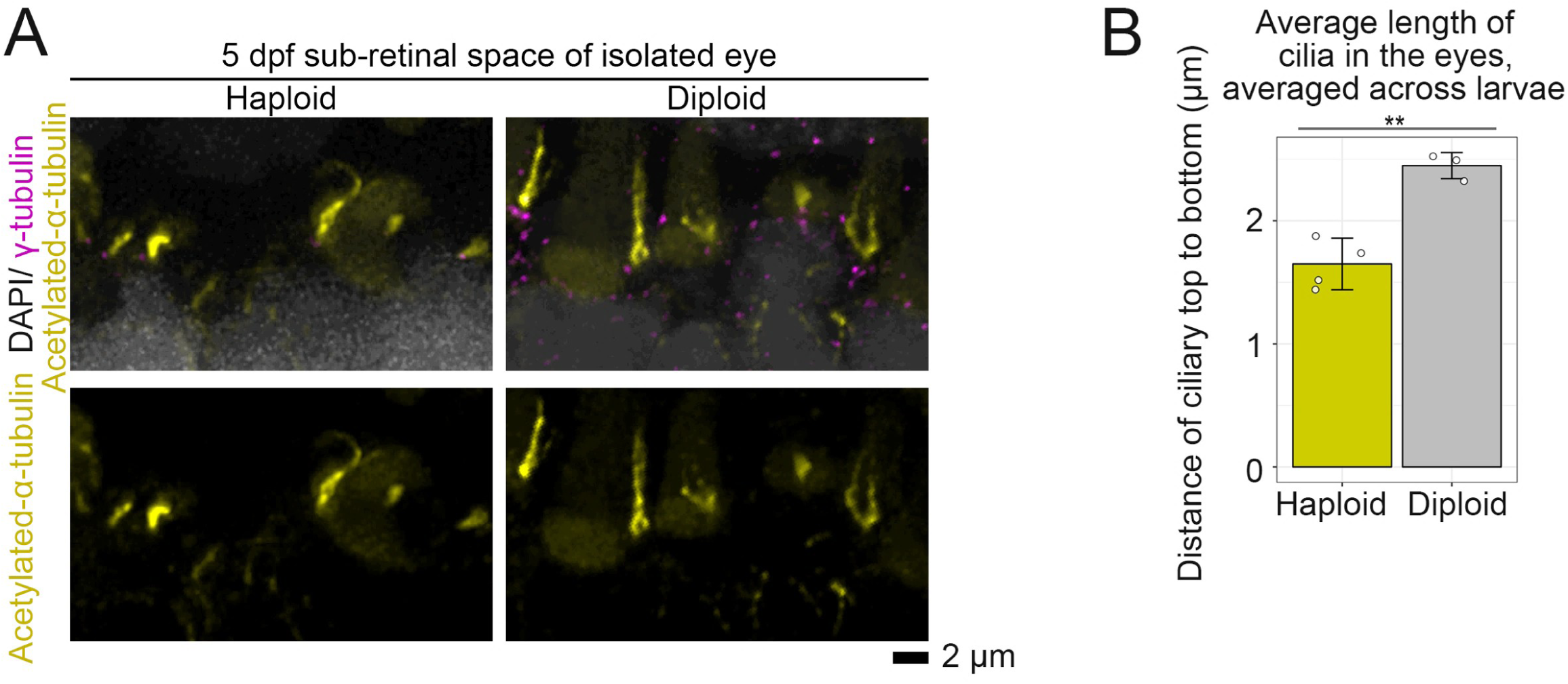
Shortened connecting cilia in haploid retinal cells. **(A)** Deconvolved immunostaining images of acetylated-α-tubulin and γ-tubulin in eyes isolated from 5-dpf haploid or diploid larvae. Scale bar: 2 μm. **(B)** Quantification of cilia length in A. Mean ± s.d. of at least 3 eyes from at least 3 larvae of 2 independent experiments, both for haploids and diploids. At least 24 cilia were analyzed per eye. **p<0.01, two-tailed t-test.

### Tissue defects in the inner retinal cell layers in haploids

The aforementioned EM images indicate drastic disorganization in the inner retinal cell layers in haploid eyes (Fig. 1). To address this haploidy-linked defect at the cellular level, we next examined the spatial distribution of retinal ganglion cells (RGCs), which mainly comprise the GCL, by immunostaining for neurolin as an RGC marker. In 2.5-dpf diploids, a continuous, even layer of RGCs formed around the lens (Fig. 4A). In contrast, 2.5-dpf haploid eyes exhibited discontinuous neurolin-positive structures with irregular thickness and jagged borders protruding toward the outer layers (Fig. 4A). The area occupied by neurolin-positive cells per total retinal area (detected by DAPI staining) was significantly reduced in haploids compared to diploids (Fig. 4B).

**Figure 4:**
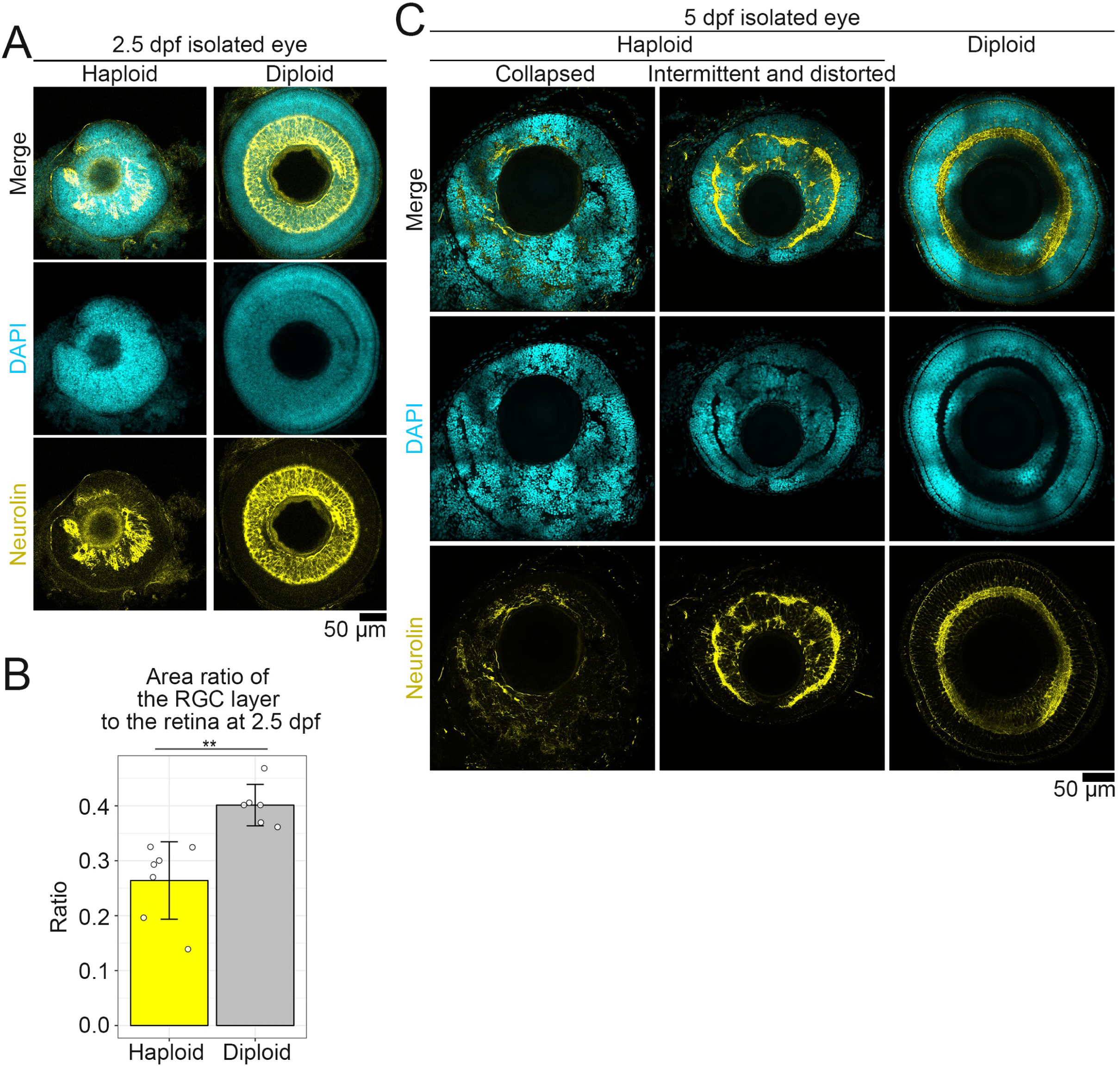
Inner tissue layer disorganization in haploid larval retina. **(A, C)** Immunostaining of neurolin in eyes isolated from haploid or diploid embryos at 2.5 dpf (A) or 5 dpf (C). DNA was stained with DAPI. Scale bar: 50 μm. **(B)** The ratio of the RGC layer area (detected by neurolin immunostaining) to the whole-retinal area (detected by DAPI staining) in A. Mean ± s.d. of at least 6 eyes from at least 6 larvae in 3 independent experiments, both for haploids and diploids. **p<0.01, two-tailed t-test. **(C)** Immunostaining of neurolin in isolated eyes from haploid or diploid larvae at 5 dpf. Scale bar: 50 μm. Representative data from at least 3 eyes from 2 independent experiments.

At later stages in diploids (5 dpf), neurolin signals showed an orderly distribution in the GCL and IPL, where RGC cell bodies and neurites are located (Laessing and Stuermer, 1996; Mumm et al., 2006), and were also observed in the INL and OPL (Fig. 4C). In 5-dpf haploids, neurolin-positive structures were still severely disorganized, with highly heterogeneous patterns among individuals (Fig. 4C). These data collectively indicate that GCL disorganization in haploids is not merely due to general developmental delay, but due to stochastic defects in the process of RGC formation and/or redistribution.

## Discussion

Haploidy has been shown to cause abnormalities in cell proliferation at the cellular level and developmental abnormalities at the organismal level in animals (Hamilton, 1966; Olbrich et al., 2017; Yaguchi et al., 2024). An important remaining question is whether haploidy-linked organismal abnormalities are explained solely by cell proliferation defects. In this study, through a systematic comparative analysis of tissue architecture in zebrafish eyes, we found that haploidy caused pleiotropic defects in tissue organization that were distinct from tissue downsizing resulting from proliferative failure.

Beyond macroscopic deformations of layered eye tissues, we observed drastic abnormalities in cellular and intracellular structures critical for visual function in haploids. More specifically, haploid eyes showed a substantial reduction in photoreceptor cells, with frequent mislocalization to the inner retinal layers. Moreover, photoreceptor cells, even when correctly located in the PCL, showed severe retardation in connecting cilia with deformed outer segments. These observations suggest that photo-sensing and signal transduction functions are profoundly impaired in haploid eyes.

Given the genome-wide transcriptional changes associated with haploidy (Menon et al., 2020; Yaguchi et al., 2024), we need to assume diverse causes for the pleiotropic tissue and cellular abnormalities in haploid organs. Meanwhile, the cellular defects in haploid eyes described above may be explained, at least in part, by loss of centrosomes or associated primary cilia. We previously found that haploid zebrafish embryos suffer progressive centrosome loss in multiple organs, including eyes, after 1 dpf (Yaguchi et al., 2024). Consistent with this finding, we observed a substantial reduction in cilia-positive cells in the outer edge of the neural retina in 5-dpf haploids (Fig. 2A and D). Interestingly, previous studies reported that zebrafish mutants with impaired ciliary function exhibit defects in photoreceptor morphogenesis or RGC layer formation (Lepanto et al., 2016; Sukumaran and Perkins, 2009; Tsujikawa and Malicki, 2004). Therefore, one possible contributor to the above-mentioned tissue defects is the previously reported centrosome instability in haploid tissues. However, the multifaceted defects described here likely reflect combined consequences of multiple haploidy-associated cellular abnormalities.

In light of haploid bioresource engineering, elucidating and manipulating haploidy-specific cellular defects has become a promising approach. For instance, we recently genetically alleviated haploidy-specific mitotic instability to achieve stable proliferation in haploid human cultured cells (Yoshizawa et al., 2026). To apply a similar framework to organismal haploid bioresources, manipulating multiple factors beyond proliferation, possibly including restoration of centrosome-cilia regulation, would be critical. Our data provide insight into diverse cellular principles underlying haploidy-linked developmental defects and clues to overcome them.

## Material and Methods

### Zebrafish

Wild-type zebrafish were obtained from the National BioResource Project Zebrafish Core Institution (NZC, Japan). Animal experiments in this study were approved by the Committee on Genetic Recombination Experiment, Hokkaido University. Fish were maintained at 28.5°C under a 14-h light and 10-h dark cycle. For in vitro fertilization, whole testes from a single male were dissected into 1 mL of cold Hank’s buffer (0.137 M NaCl, 5.4 mM KCl, 0.25 mM Na2HPO4, 0.44 mM KH2PO4, 1.3 mM CaCl2, 1.0 mM MgSO4 and 4.2 mM NaHCO3) to collect sperm. Sperm DNA was inactivated by irradiating with 254 nm UV light (LUV-6, AS ONE) at a distance of 30 cm for 1 min with gentle pipetting every 20 s. Eggs were extruded from females immobilized by anaesthesia (A5040, Sigma-Aldrich; 0.08% ethyl 3-aminobenzoate methanesulfonate salt, pH 7.2). For insemination, we added 200 μl sperm solution to approximately 100 eggs. After the chorion inflation, embryos were grown in Embryo medium at 28.5°C until use.

To inhibit pigmentation, we treated embryos with 0.03 g l^−1^ N-phenylthiourea (P7629, Sigma-Aldrich) at 0.5 or 1 dpf. Dechorionation was performed by treating embryos with 0.2–5 mg ml^−1^ pronase (11459643001, Roche) in Embryo medium for 1–2 min at 25°C, followed by washing with Embryo medium. The remaining chorion was manually removed.

### Electron microscopy

Larvae were fixed at 5 dpf by immersion in fixative solution (2 % paraformaldehyde, 2.5 % glutaraldehyde in PBS). Larvae were fixed at 25°C during the first 30 minutes, and then incubated overnight at 4°C. Eyes were isolated from the fixed larvae, followed by removal of the sclera using forceps in phosphate-buffered saline (PBS) (Zhang and Leung, 2010). To preliminarily specify the distributions of photoreceptor cells within the specimen, the isolated eyes were stained with 4′,6-Diamidino-2-phenylindole, dihydrochloride (DAPI) and lectin PNA from Arachis hypogaea (peanut), Alexa Fluor™ 568 conjugate (PNA) (see below) overnight, mounted in 5% low-melting-point agarose (PrimeGel™ Agarose LMT 1-20K; Takara Bio, 5805A) in PBS and filmed under NIKON A1 Rsi microscope. The mounted eyes were stored in fixative solution at 4 °C until use.

The mounted eyes were washed with 0.1 M cacodylate buffer (pH 7.3) three times, further fixed with 1.5% potassium ferrocyanide and 1% osmium tetroxide in the same buffer at 4 °C for 1 h, and then washed five times with Milli-Q water. After the samples were immersed in 0.5% uranyl acetate in Milli-Q water for 2 h, they were dehydrated with an ethanol series (30%, 70%, 85%, 95%, and 99.5% for 15 min each and three times in 100% for 20 min) and propylene oxide for 20 min twice, followed by infiltration with Epon 812 resin (TAAB). Polymerization of the resin was performed at 65°C for 72 h. Sections (200 nm) were cut with a diamond knife on an ultramicrotome (Leica EM UC6, Leica Microsystems) and placed on a piece of silicon wafer. The sections were stained with uranyl acetate and lead nitrate, then observed with a field-emission scanning electron microscope JSM-IT800SHL (JEOL) at 3 kV using a back-scattered electron detector.

### Immunofluorescence staining

Larvae were fixed with 4% paraformaldehyde in PBS for at least 4 h at 28.5°C or overnight at 4℃, followed by partial digestion with a cold trypsin mixture for 3 min. Fixed larvae were manually deyolked in 0.1% Triton X-100 in PBS and permeabilized with 0.5 or 1% Triton X-100 in PBS overnight at 4°C, followed by treatment with bovine serum albumin (BSA) blocking buffer (150 mM NaCl, 10 mM Tris-HCl, pH 7.5, 5% BSA and 0.1% Tween 20) for >30 min at 4°C or for 2 h at 28.5°C. Larvae were subsequently incubated with primary antibodies for >24 h at 4°C and with secondary antibodies overnight at 4°C. Following each antibody incubation, larvae were washed three times with 0.1% Triton X-100 in PBS. Acetylated-α-tubulin or F-actin was poststained by incubating larvae with fluorescence-conjugated specific antibody (see below) for >24 h at 4°C or with Acti-stain 555 phalloidin for >4 h at 25°C, respectively, followed by wash with 0.1% Triton X-100 in PBS. Immunostained larvae were mounted in Fluoromount (K024, Diagnostic BioSystems), RapiClear 1.52 (RC152001, SunJin Lab), or 80% glycerol in PBS.

### Antibodies and dyes

Antibodies were purchased from suppliers and used at the following dilutions: mouse anti-arrestin3 (Abcam, ZPR-1, ab174435; 1:400-500); mouse anti-gamma-tubulin (Abcam, GTU-88, ab11316; 1:500); mouse anti-neurolin antibody (DSHB, zn-8; 1:200); Alexa Fluor 488-conjugate rabbit monoclonal anti-acetylated-α-tubulin (Cell Signaling Technology, D20G3; 1:200) and fluorescence-conjugated secondaries (Alexa Fluor 594 or Alexa Fluor 647; Jackson ImmunoResearch Laboratories, Inc. or Abcam).

Fluorescent dyes and probes were purchased from suppliers and used at the following dilutions: DAPI (Dojindo, D523; 1:50–800); Alexa Fluor 568-conjugated PNA (Invitrogen, L32458; 1:500-800); and Acti-stain 555 phalloidin (Cytoskeleton, Inc., PHDH1; 1:20).

### Microscopy

Fixed larval eyes were observed under either of the following microscopes: an A1Rsi microscope (Nikon) equipped with a ×40 1.30 NA or ×100 1.40 NA apochromatic oil-immersion objective lens, an LU-N4S laser unit (403, 489, 562, and 639 nm), and a GaAsP detector unit; an A1Rsi microscope (Nikon) equipped with a ×100 1.35 NA apochromatic silicon oil-immersion objective lens, an LU-N4S laser unit (405, 488, 561, and 640 nm), and an A1-DUG detector unit; an A1 microscope (Nikon) equipped with a ×100 1.40 NA apochromatic oil-immersion objective lens and an LU4 laser unit (403, 487, 562, and 636 nm); or an AX/AX R microscope (Nikon) equipped with a ×100 1.45 NA apochromatic oil-immersion objective lens, an LUA-S6 laser unit (405, 488, 561, and 640 nm), and an NSPARC detector unit. Image acquisition was controlled using NIS-Elements (Nikon). Image acquisition was controlled using NIS-Elements (Nikon). Deconvolution was performed for cilia panels in the figures and analysis using the Automatic algorithm in NIS-Elements (Nikon).

### Statistical analysis

Analyses for significant differences between the two groups were conducted using a two-tailed Student’s t-test in Excel (Microsoft). We set statistical significance at p<0.05. p-values are indicated in figures or the corresponding figure legends.

---

**Figure S1.**
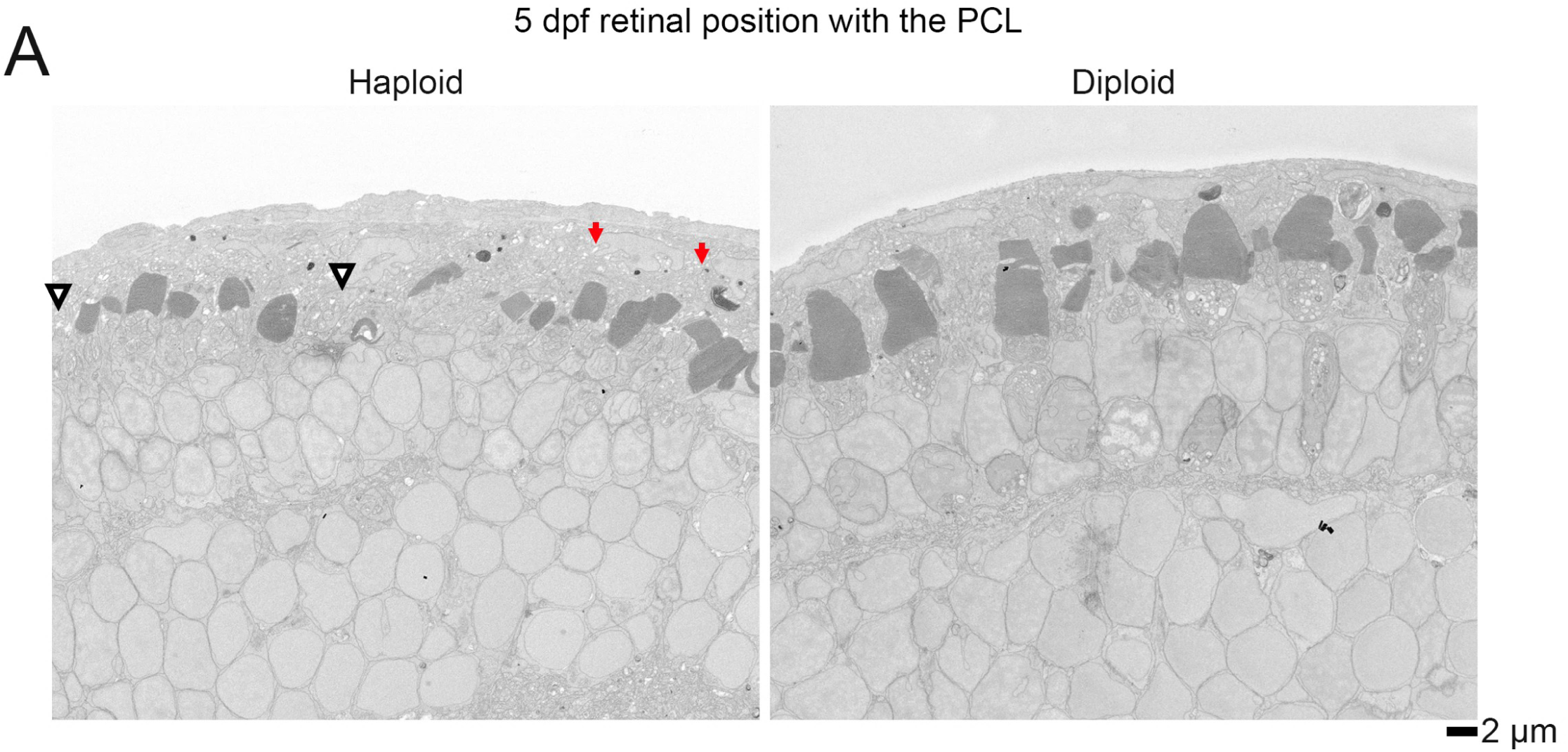
**(A)** Higher magnification EM images of retinal layers in eyes isolated from 5-dpf haploid or diploid larvae. The images were scanned at 2000× magnification. Arrowheads indicate the PCL area devoid of outer segments of photoreceptor cells. Arrows indicate positions where the layer structure is disrupted, yet the outer segments of photoreceptor cells remain. Scale bar: 2 μm. Representative data from 3 eyes from 2 independent experiments.

**Supplemental Video S1**

Three-dimensional projection of Arrestin3 and acetylated-α-tubulin immunostaining with fluorescence-labeled PNA co-staining in 5-dpf haploid eyes. Representative data from 4 eyes from 2 independent experiments. The image is shown at 24 pixel/μm.

**Supplemental Video S2**

Three-dimensional projection of Arrestin3 and acetylated-α-tubulin immunostaining with fluorescence-labeled PNA co-staining in 5-dpf diploid eyes. Representative data from 4 eyes from 2 independent experiments. The image is shown at 24 pixel/μm.

## Acknowledgment

We are grateful to Sreelaja Nair and Tomoya Kotani for valuable discussions, to Kentaro Kobayashi and other members of the Nikon Imaging Center at Hokkaido University for imaging technical support, and to NBRP for providing the RIKEN WT zebrafish strain. This work was supported by COI-NEXT Support Unit for Imaging Science at Kento. DS are supported by JST SPRING, Grant Number JPMJSP2119. This work was supported by JSPS KAKENHI (Grant Numbers JP19KK0181, JP19H05413, JP19H03219, JPJSBP120193801, JP21K19244, JP22H04926, JP24K21956, JP24KK0139, and JP24K02017 to R.U.), the Princess Takamatsu Cancer Research Fund, the Kato Memorial Bioscience Foundation, the Orange Foundation, the Smoking Research Foundation, Daiichi Sankyo Foundation of Life Science, the Akiyama Life Science Foundation, the Hoansha Foundation, Sumitomo Electric Group CSR Foundation, and the Terumo Life Science Foundation to R.U.

## Author Contributions

Conceptualization, D.S., K.Y., and R.U.; Methodology, D.S., A.S., K.Y., Y.Y., K.N., S.Y., and R.U.; Investigation, D.S., A.S., K.Y., R.S. and S.Y.; Formal Analysis, D.S., and R.S.; Resources, D.S., A.S., K.Y., R.S., S.Y., and R.U.; Supervision, R.U.; Writing – Original Draft, D.S., and R.U.; Writing – Review & Editing, D.S., S.Y., and R.U.; Funding Acquisition, D.S., S.Y., and R.U.

## Data and resource availability

All relevant data and details of resources can be found within the article and its supplementary information.

## Reference

Amini, R., Rocha-Martins, M. and Norden, C. (2018). Neuronal Migration and Lamination in the Vertebrate Retina. Front. Neurosci. 11, 742.

Amini, R., Bhatnagar, A., Schlüßler, R., Möllmert, S., Guck, J. and Norden, C. (2022). Amoeboid-like migration ensures correct horizontal cell layer formation in the developing vertebrate retina. eLife 11, e76408.

Beryozkin, A., Matsevich, C., Obolensky, A., Kostic, C., Arsenijevic, Y., Wolfrum, U., Banin, E. and Sharon, D. (2021). A new mouse model for retinal degeneration due to Fam161a deficiency. Sci. Rep. 11, 2030.

Di Minin, G., Rüegg, A. B., Halter, K., Fuhrer, T., Kleele, T., Zamboni, N. and Wutz, A. (2026). Mitochondrial metabolic imbalance drives diploidization in mouse haploid embryonic stem cells via NADPH overload. Nat. Commun. 17, 4359.

Edwards, M. M., Zuccaro, M. V., Sagi, I., Ding, Q., Vershkov, D., Benvenisty, N., Egli, D. and Koren, A. (2021). Delayed DNA replication in haploid human embryonic stem cells. Genome Res. 31, 2155–2169.

Fadool, J. and Dowling, J. (2008). Zebrafish: A model system for the study of eye genetics. Prog. Retin. Eye Res. 27, 89–110.

Hamilton, L. (1966). The role of the genome in the development of the haploid syndrome in Anura. Development 16, 559–568.

Hanke-Gogokhia, C., Chiodo, V. A., Hauswirth, W. W. and Frederick, J. M. (2018). Rescue of cone function in cone-only Nphp5 knockout mouse model with Leber congenital amaurosis phenotype. Mol. Vis. 24, 834–846.

Ishida-Ishihara, S., Yaguchi, K., Miura, S., Nomura, R., Wang, Q., Yoshizawa, K., Sato, K., Yang, G., Veszelyi, K., Banhegyi, G., et al. (2024). Fragility of ER homeostatic regulation underlies haploid instability in human somatic cells. J. Biol. Chem. 300, 107909.

Komen, H. and Thorgaard, G. H. (2007). Androgenesis, gynogenesis and the production of clones in fishes: A review. Aquaculture 269, 150–173.

Krock, B. L., Mills-Henry, I. and Perkins, B. D. (2009). Retrograde Intraflagellar Transport by Cytoplasmic Dynein-2 Is Required for Outer Segment Extension in Vertebrate Photoreceptors but Not Arrestin Translocation. Investig. Opthalmology Vis. Sci. 50, 5463.

Laessing, U. and Stuermer, C. A. O. (1996). Spatiotemporal pattern of retinal ganglion cell differentiation revealed by the expression of neurolin in embryonic zebrafish. J. Neurobiol. 29, 65–74.

Lepanto, P., Davison, C., Casanova, G., Badano, J. L. and Zolessi, F. R. (2016). Characterization of primary cilia during the differentiation of retinal ganglion cells in the zebrafish. Neural Develop. 11, 10.

Lessieur, E. M., Fogerty, J., Gaivin, R. J., Song, P. and Perkins, B. D. (2017). The Ciliopathy Gene *ahi1* Is Required for Zebrafish Cone Photoreceptor Outer Segment Morphogenesis and Survival. Investig. Opthalmology Vis. Sci. 58, 448.

Luo, C. and Li, B. (2003). Diploid-dependent regulation of gene expression: a genetic cause of abnormal development in fish haploid embryos. Heredity 90, 405–409.

Menon, T., Borbora, A. S., Kumar, R. and Nair, S. (2020). Dynamic optima in cell sizes during early development enable normal gastrulation in zebrafish embryos. Dev. Biol. 468, 26–40.

Mumm, J. S., Williams, P. R., Godinho, L., Koerber, A., Pittman, A. J., Roeser, T., Chien, C.-B., Baier, H. and Wong, R. O. L. (2006). In Vivo Imaging Reveals Dendritic Targeting of Laminated Afferents by Zebrafish Retinal Ganglion Cells. Neuron 52, 609–621.

Norden, C. (2023). A Fish Eye View: Retinal Morphogenesis from Optic Cup to Neuronal Lamination. Annu. Rev. Cell Dev. Biol. 39, 175–196.

Ocalewicz, K. (2024). Quality of fish eggs and production of androgenetic and gynogenetic doubled haploids (DHs). Fish Physiol. Biochem. 50, 1947–1957.

Ojeda Naharros, I., Cristian, F. B., Zang, J., Gesemann, M., Ingham, P. W., Neuhauss, S. C. F. and Bachmann-Gagescu, R. (2018). The ciliopathy protein TALPID3/KIAA0586 acts upstream of Rab8 activation in zebrafish photoreceptor outer segment formation and maintenance. Sci. Rep. 8, 2211.

Olbrich, T., Mayor-Ruiz, C., Vega-Sendino, M., Gomez, C., Ortega, S., Ruiz, S. and Fernandez-Capetillo, O. (2017). A p53-dependent response limits the viability of mammalian haploid cells. Proc. Natl. Acad. Sci. 114, 9367–9372.

Purdom, C. E. (1969). Radiation-induced gynogenesis and androgenesis in fish. Heredity 24, 431–444.

Richardson, R., Tracey-White, D., Webster, A. and Moosajee, M. (2017). The zebrafish eye—a paradigm for investigating human ocular genetics. Eye 31, 68–86.

Schmitt, E. A. and Dowling, J. E. (1999). Early retinal development in the zebrafish,Danio rerio: Light and electron microscopic analyses. J. Comp. Neurol. 404, 515–536.

Shi, Y., Obert, E., Rahman, B., Rohrer, B. and Lobo, G. P. (2017). The Retinol Binding Protein Receptor 2 (Rbpr2) is required for Photoreceptor Outer Segment Morphogenesis and Visual Function in Zebrafish. Sci. Rep. 7, 16207.

Sukumaran, S. and Perkins, B. D. (2009). Early defects in photoreceptor outer segment morphogenesis in zebrafish ift57, ift88 and ift172 Intraflagellar Transport mutants. Vision Res. 49, 479–489.

Tsujikawa, M. and Malicki, J. (2004). Intraflagellar Transport Genes Are Essential for Differentiation and Survival of Vertebrate Sensory Neurons. Neuron 42, 703–716.

Wutz, A. (2014). Haploid animal cells. Development 141, 1423–1426.

Yaguchi, K., Yamamoto, T., Matsui, R., Tsukada, Y., Shibanuma, A., Kamimura, K., Koda, T. and Uehara, R. (2018). Uncoordinated centrosome cycle underlies the instability of non-diploid somatic cells in mammals. J. Cell Biol. 217, 2463–2483.

Yaguchi, K., Saito, D., Menon, T., Matsura, A., Hosono, M., Mizutani, T., Kotani, T., Nair, S. and Uehara, R. (2024). Haploidy-linked cell proliferation defects limit larval growth in zebrafish. Open Biol. 14, 240126.

Yoshizawa, K., Yaguchi, K. and Uehara, R. (2020). Uncoupling of DNA Replication and Centrosome Duplication Cycles Is a Primary Cause of Haploid Instability in Mammalian Somatic Cells. Front. Cell Dev. Biol. 8, 721.

Yoshizawa, K., Raj Singh, H., Kathirvel, P., Zhu, J. and Uehara, R. (2026). Cep192 insufficiency underlies haploid instability in human cells. bioRxiv doi: 10.64898/2026.03.18.712690

Zhang, L. and Leung, Y. F. (2010). Microdissection of Zebrafish Embryonic Eye Tissues. J. Vis. Exp. 2028.

